# L-type voltage-gated calcium channel regulation of *in vitro* human cortical neuronal networks

**DOI:** 10.1101/464677

**Authors:** William Plumbly, Nicholas J. Brandon, Tarek Z. Deeb, Jeremy Hall, Adrian J. Harwood

**Affiliations:** Neuroscience and Mental Health Research Institute, Cardiff University, Cardiff, CF24 4HQ, UK; Neuroscience, IMED Biotech Unit, AstraZeneca, 35 Gatehouse Dr, Waltham, MA 02451, USA; Department of Neuroscience, Tufts University School of Medicine, Boston, MA, USA

## Abstract

The combination of *in vitro* multi-electrode arrays (MEAs) and the neuronal differentiation of stem cells offers the capability to study human neuronal networks from patient or engineered human cell lines. Here, we use MEA-based assays to probe synaptic function and network interactions of hiPSC-derived neurons. Neuronal network behaviour first emerges at approximately 30 days of culture and is driven by glutamate neurotransmission. Over a further 30 days, inhibitory GABergic signalling shapes network behaviour into a synchronous regular pattern of burst firing activity and low activity periods. Gene mutations in L-type voltage gated calcium channel subunit genes are strongly implicated as genetic risk factors for the development of schizophrenia and bipolar disorder. We find that, although basal neuronal firing rate is unaffected, there is a dose-dependent effect of L-type voltage gated calcium channel inhibitors on synchronous firing patterns of our hiPSC-derived neural networks. This demonstrates that MEA assays have sufficient sensitivity to detect changes in patterns of neuronal interaction that may arise from hypo-function of psychiatric risk genes. Our study highlights the utility of *in vitro* MEA based platforms for the study of hiPSC neural network activity and their potential use in novel compound screening.

## Introduction

Understanding how neurons interact both within local networks or as components of neural circuits is a key objective for the investigation of brain function throughout development and in neuropsychiatric disorders. Study of neural interactions in animals *in vivo*, in intact brain slices or in isolated primary neurons in culture has been a mainstay of neuroscience for more than a century. This has been augmented by human studies investigating functional (EEG and MEG) and structural (MRI) connectivity. However, the study of the development and function of human neural networks at the cellular level has been more difficult, largely due to the difficulty of combining cell culture methods and longitudinal electrophysiological experimentation. This is now changing with the opportunity to develop neurons derived from human induced pluripotent stem cells (hiPSC) and the advent of user-friendly multi-electrode array (MEA) systems. These offer the potential to generate small-scale human neural networks within a familiar context and investigate their development and function *in vitro* ^1^.

Dissociated primary rodent neurons *in vitro* self-organise to form functional networks ^2-4^, which spontaneously exhibit coordinated action potential bursts (synchronised bursts; SBs) that can be detected across spatially separated electrodes of an MEA ^5,6^. This culture-wide, synchronised bursting behaviour correlates with an increased basal firing rate and emerges after an approximate two-week period of asynchronous firing. Recently, it has been shown that this synchronous behaviour may develop further to form a temporal pattern of regular low and high activity periods lasting for tens of seconds ^7^, behaviour which is sensitive to pharmacological manipulation and in particular AMPA receptor activity. MEA methods to study neuronal networks derived from hiPSC are less established. However, a number of MEA studies have shown that human stem cell-derived neurons develop spontaneous activity ^8^, which increases during an extended period of development ^9-12^. With prolonged incubation, the emergence of short SBs across multiple electrodes of the array has been observed ^10^. These studies have established the use of MEAs for pharmacological profiling and toxicology testing ^8,10,13^. However, it is currently not known whether such methods have the discriminatory sensitivity to investigate the effects of genetic risk in patient hiPSCs derived cultures, thereby offering a platform for drug development.

In this study, we demonstrate the use of MEA assays to probe synaptic function and functional neural networks in hiPSC-derived neuronal cultures. We firstly investigated the development of networked behaviour in *in vitro* neuronal cultures on MEAs and pharmacologically characterised network behaviour. We then use hiPSC-derived MEAs to probe the effects of L-type voltage-gated calcium channel activity on network behaviour. These channels have been implicated in several aspects of network function *in vivo*, including the synchronisation of calcium events ^14,15^ and the regulation of hippocampal oscillatory activity ^16,17^. Significantly, genetic variants in *CACNA1C* and *CACNA1D* (coding for the alpha-1 subunits Ca_v_ 1.2 and Ca_v_ 1.3 respectively) are some of the most robust genetic risk alleles associated with the development of mental health disorders, including schizophrenia and bipolar disorder. By use of selective L-type channel inhibitors we demonstrate a specific modulation of SB behaviour in hiPSC-derived neural networks without changes in basal neuronal activity. These results demonstrate the potential of hiPSC-based MEA assays to detect functional changes arising from genetic risk loci for psychiatric disorders.

## Results

### Creation of hiPSC-derived neuronal networks

Neurons were differentiated from human iPSC by a dual-SMAD monolayer protocol and re-plated on MEAs at 30DIV. MEA cultures were maintained in astrocyte conditioned medium (ACM) and in a 2% O_2_atmosphere to enhance neuronal maturation ^18,19^. Parallel cell cultures to those on MEAs were examined by immunocytochemistry at 50 days post re-plating (50DPP; corresponding to 50 days after cells were re-plated on MEAs or 80 days of total differentiation; Figure 1). All imaged neurons expressed the neuron-specific microtubule associated protein MAP2, of which 64 ± 9.0 % co-expressed the vesicular glutamate transporter vGLUT1 and 3.6 ± 1.2% of cells expressed the GABA synthesising enzyme GAD67 (Figure 1D), indicating a mixed culture of predominately glutamatergic neurons with a small population of GABAergic cells. We also found that 73.5 ± 9.7% of nuclei expressed CTIP2 with 37.7 ± 8.4% also expressing SATB2 (Figure 1B), suggesting a deep-layer cortical identity of most neurons primarily corresponding to layers 6 and 5a (Figure 1D). Higher magnification images revealed that neurons also expressed the key excitatory synaptic scaffolding protein PSD95 and the obligatory glycine-binding NMDA receptor subunit, GluN1, suggesting the presence of excitatory synapses (Figure 1C).

**Figure 1.**
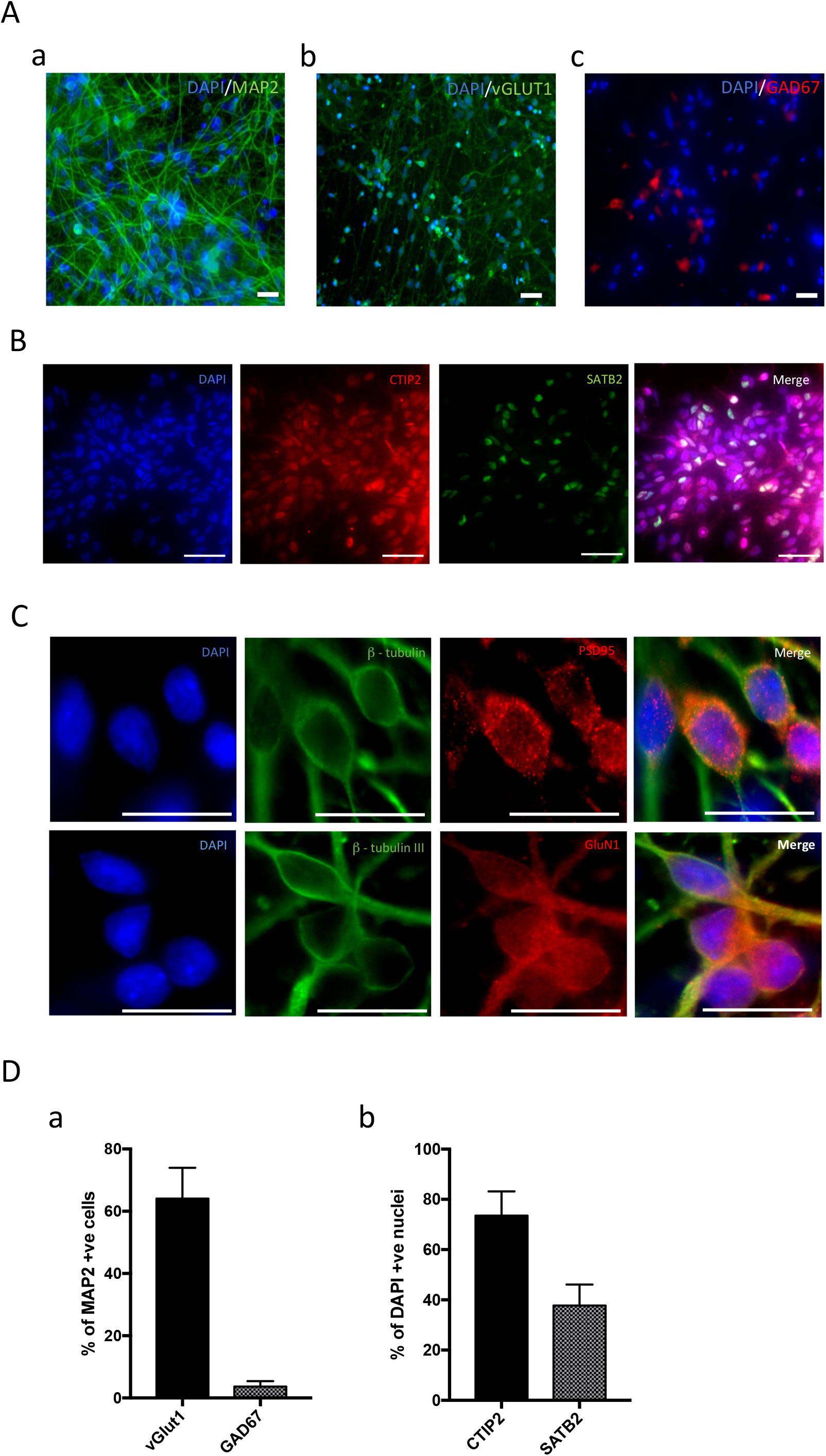
Morphology of hiPSC derived neurons. (A) Fluorescent images showing identity of differentiated neurons at 50DPP. Neurons show expression of MAP2 (a), with a majority of these also expressing vGLUT1 (b) and a minority expressing GAD67 (c). (B) Layer marker expression in neurons at 50DPP. (C) Expression of excitatory post-synaptic markers in neurons at 50DPP. (D) Summary plots showing quantification of markers highlighting (a) neuronal fate (b) cortical layer markers. Scale bars in A, B & C = 50 μm. Plots in D show means + SD.

In order to confirm the functional maturation of the hiPSC derived neuron, we performed single-cell patch clamp experiments (Figure 2A), which provides key insights into the basal physiological state of the neurons. Resting membrane potential (V_m_) and input resistance of neurons significantly reduced throughout differentiation (−30.6 ± 10.2 mV to −44.8 ± 10.0 mV and 1.26 ± 0.66 GΩ to 0.92 ± 0.41 GΩ respectively), indicating that neurons were developing over time, although this was not reflected in any change to the membrane time constant (tau) (Figure 2C). Patched neurons were subsequently classified according to their ability to fire induced and spontaneous action potentials (iAP and sAPs respectively). Cells were held at −70mV via constant current injection and iAPs were elicited by injecting additional depolarising current steps. Neurons were subsequently coded based upon their voltage response. At 50DPP, 83.9% of neurons were classed as active, which included neurons showing single iAPs, attempts at trains or full iAP trains (Figure 2A; see methods). The AP threshold and average event amplitude of these iAPs was unchanged over development, however neurons did exhibit AP with faster rise times and shorter half-widths at 50DPP compared to 30DPP (Figure 2C), highlighting the maturation of the cells over these 20 days. Finally, at 50DPP, 22.9% of patched neurons exhibited sAPs, as determined by continuous recording of cells at their resting membrane potential (I = 0; Figure 2B).

**Figure 2.**
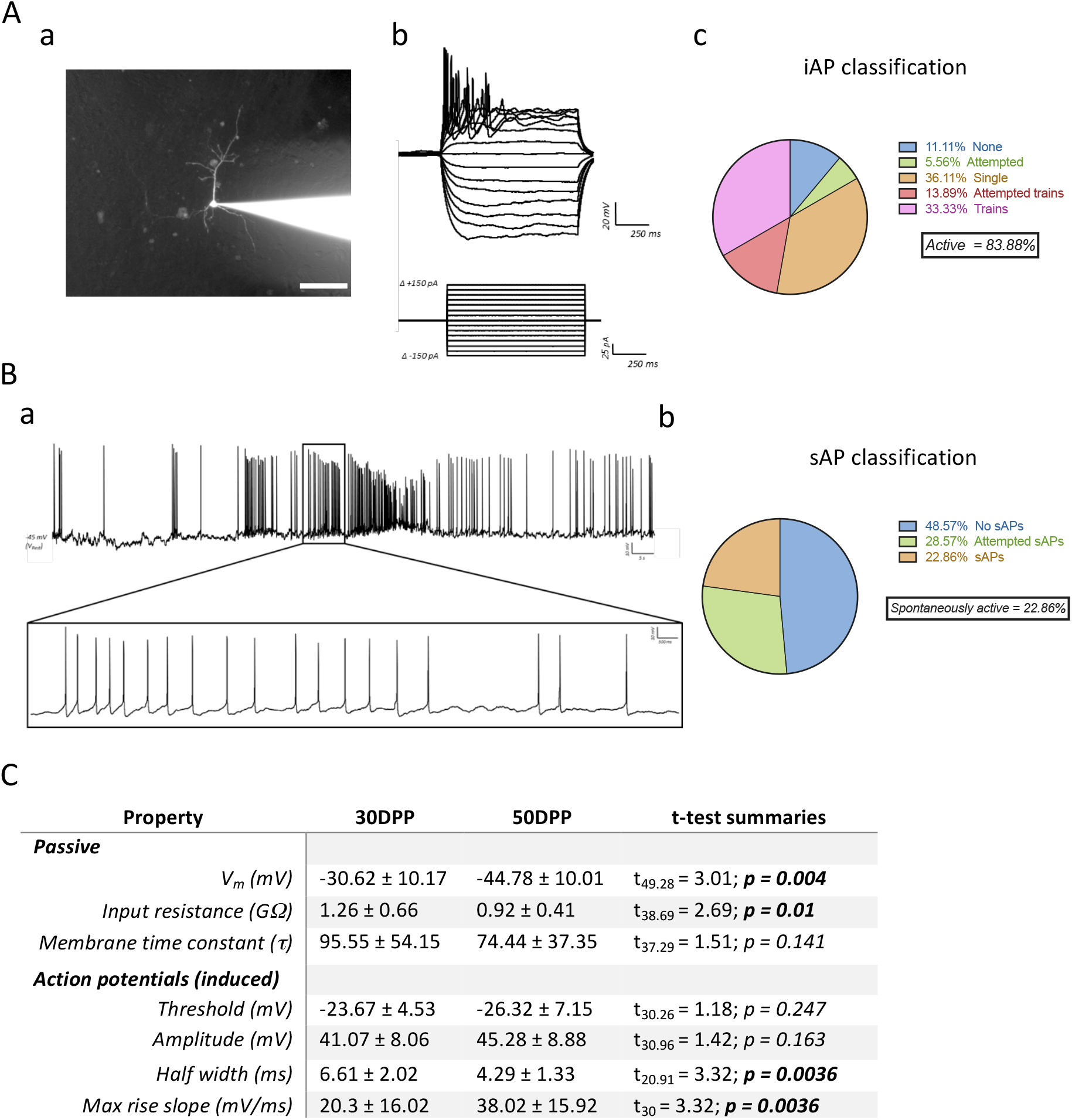
Intrinsic and action potential properties of hiPSC derived cortical neurons. (A) Induced action potential (iAP) formation in neurons. (a) Representative fluorescent image of a hiPSC derived neuron during single cell patching filled with Alexa Fluor 488 dye, highlighting pyramidal-like morphology. (b) Current injection protocol used for determination of passive and action potential properties of patched neurons. Cells were held at −70 mV. (c) Classification of patched neurons based upon their induced action potential response. Full action potentials had an overshoot of > 0 mV; trains contained ≥ 2 full action potentials. (B) Spontaneous action potentials (sAP) in hiPSC derived neurons. (a) Voltage trace showing presence of spontaneous activity in one patched cell. Inset highlights period of higher-frequency firing. (b) Classification of patched cells based upon spontaneous action potential formation. Valid action potentials had overshoots of > 0 mV. (C) Table summarising the intrinsic and action potential properties of patched iPS cell derived neurons. N = 29 (30DPP), 36 (50DPP).

Our single cell electrophysiology and imaging experiments indicated that our protocols yielded mature and functionally active neurons. To examine the development of networks in the cultures, differentiated hiPSC-derived neurons were plated as drop cultures onto MEAs possessing an 8×8 grid of substrate embedded electrodes, spaced 200 μm apart. Spike detection from filtered raw voltage recording made from each electrode utilised the threshold-based method of QuianQuiroga and co-workers ^20^ and following quality control spikes were timestamped for subsequent analysis. Spontaneous activity of cultures changed markedly over the 60 days of recorded development (Figure 3A). Initial activity was very low with a mean firing rate of 0.065 ± 0.021 Hz and no detectable spike bursts, but increased to a firing rate of 0.75 ± 0.21 Hz by 20DPP with 232 ± 103 bursts. The most substantial increase in activity occurred between 20-30 DPP with a 2-3-fold increase in spike rate and burst number, while after 30 DPP basal activity levels remain largely stable. These results are consistent with an increase in excitability previously seen throughout hiPSC-derived culture development ^8-11^.

**Figure 3.**
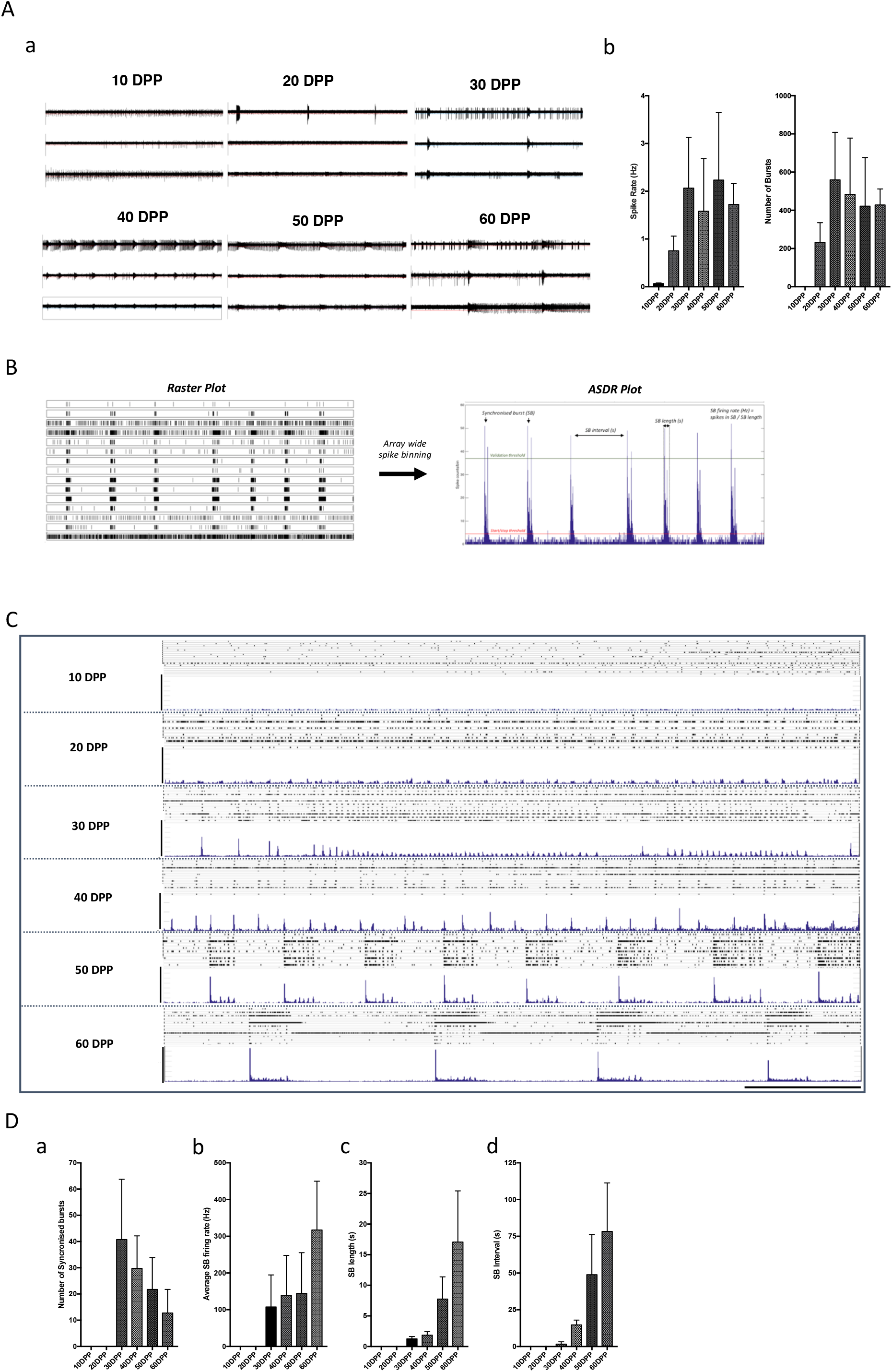
Changes in population activity of hiPSC derived neurons recorded with multi electrode arrays (MEAs). (**A**) (a) Representative voltage traces from three electrodes of the same MEA culture showing the changes in activity over a period of 60 days (DPP = days post plating). (b) Basal excitability measured over culture development showing mean spike rate and number of single unit bursts detected. (**B**) Conversion of time-stamp raster data to array wide spike detection rate (ASDR) plots, which form the basis of synchronised burst (SB) analyses. Plots are created by counting spikes in each 200 ms bin for each electrode, summing the counts for each bin and serially plotting the results. ASDR plots clearly show the occurrence of a culture-wide SB and allow the calculation of SB firing rates, lengths and intervals. (**C**) Development of array wide neuronal activity over 60 days of culture. For each time point, upper panel shows raster plot and lower panel shows ASDR plot. Horizontal scale bar = 100 seconds; vertical scale bars = 100 spikes per 200ms bin (**D**) Changes in synchronised burst (SB) activity over culture development showing (a) number of SBs detected over recording period (10 minutes); (b) mean firing rate of SBs – calculated as the number of spikes in individual SBs / length of that burst; (c) mean length of SBs and (d) mean interval between SBs. All summary data show means + standard deviations.

The emergence of synchronised bursting (SB) coincided with the switch to high neuronal activity levels seen at 30 DPP (Figure 3A). This change in behaviour can be seen more clearly in raster and array – wide spike detection rate (ASDR) plots (Figure 3B & C). At 30DPP, SBs were short in length (1.25 ± 0.37 s) with a firing rate of 107.60 ± 87.06 Hz (Figure 3D). Over the next 30 days, the number of SBs *decreased*, with a concomitant increase in both the length of the bursts and the interval between them (17.07 ± 8.34 s and 78.33 ± 33.00 s respectively at 60DPP). Although SB firing is present from 30DPP, there was a clear shift in the pattern of array– wide activity from 50 DPP into a regular pattern of more active periods (MAPs) and less active periods (LAPs) each lasting >30 seconds. Once networks have entered this state, the MAP/LAP pattern evolves further by increasing in MAP (SB) and LAP (interval) lengths (Figure 3D). Finally, it should be noted that the changes in synchronised activity patterns occur with little change of overall basal neuronal activity (Figure 3A).

### Pharmacological profiling of hiPSC derived neuronal networks

To establish whether the extended SB firing patterns arise via synaptic activity, we probed our MEA cultures with pharmacological agents that alter glutamate/GABA signalling. At 20 DPP, when no SB activity is seen, neuronal cultures are unresponsive to AMPA, NMDA and GABA_A_ receptor inhibition, but, interestingly, sensitive to 10μM GABA, which strongly attenuated spontaneous activity (Figure S1). At the later 50 DPP stage when sustained SB firing has fully matured, inhibition of glutamate signalling by application of either the AMPA receptor antagonist CNQX and or the NMDA receptor antagonist APV completely suppressed the SBF pattern (Figure 4A, B & C). In both cases, SB patterns rapidly re-establish after drug washout. Importantly, neither drug had a substantial effect on the basal rate of neuronal firing (Figure 5C), indicating that both AMPA and NMDA receptor mediated neurotransmission is required for network synchrony, not single cell firing.

**Figure 4.**
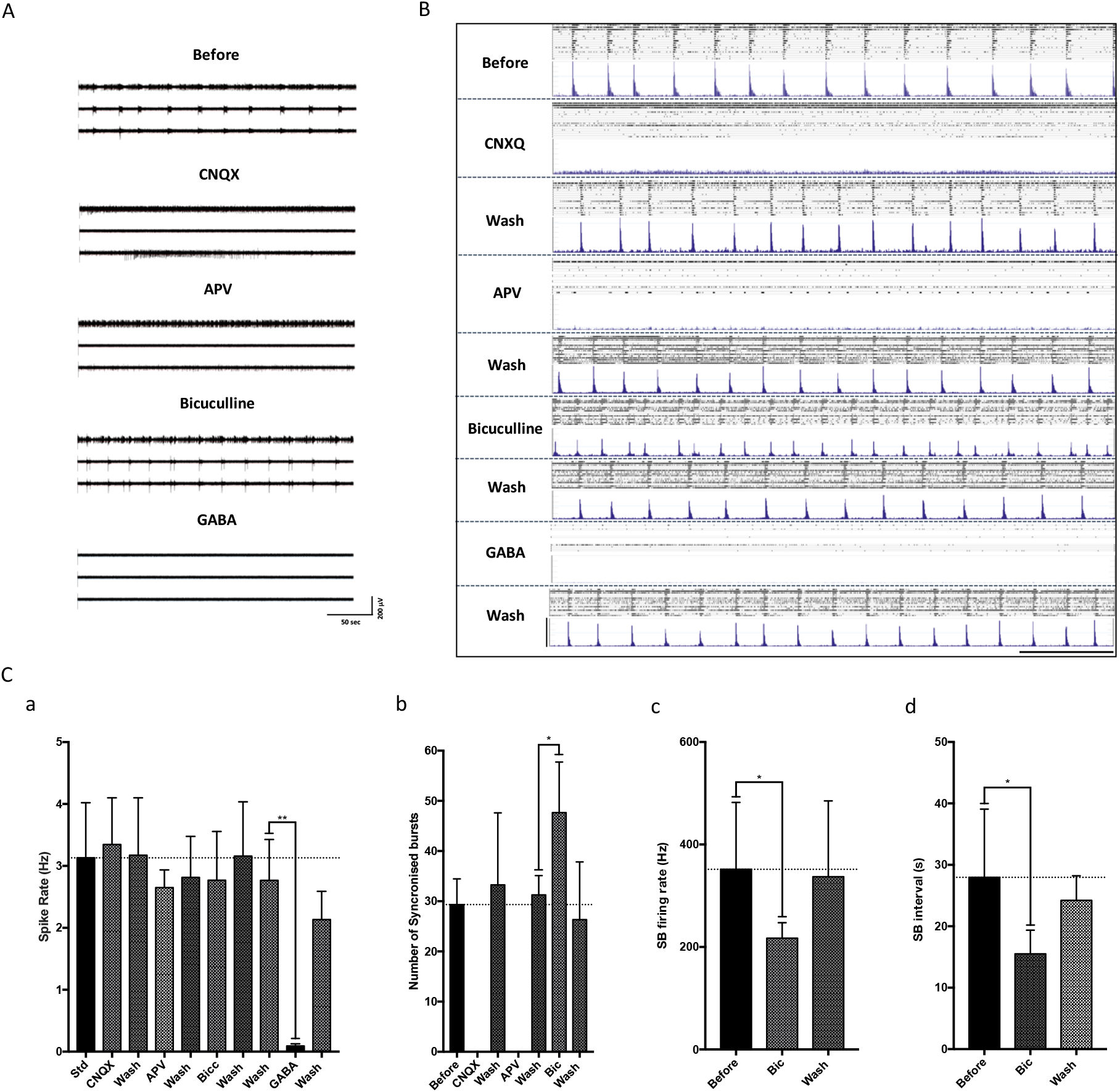
Pharmacological profiling of the network activity of hiPSC derived cortical neurons. (A) Filtered voltage traces of the same three electrodes recording the same MEA culture during acute exposure to NMDA, AMPA and GABA_A_ receptor inhibitors. (B) Raster (upper panel) and ASDR (lower panel) plots of recordings of the same MEA showing the culture – wide response to acute exposure of CNQX (50 μM), AMPA (50 μM), bicuculline (10 μM) and GABA (10 μM). Established network activity in the cultures are dependant on both AMPA and NMDA receptor activity, while the coordinated behaviour is regulated by GABAergic tone. (C) Summary bar plots showing the response of MEA cultures to acute pharmacological treatment: (a) basal spike rate following all drug treatments and washes; (b) number of synchronised bursts; synchronised burst firing rate (c) and interval (d) following bicuculline exposure. All plots show means + SD. * = p < 0.05, ** = p < 0.01 following paired t – tests. Number of arrays = 9, all at 50-55DPP.

**Figure 5.**
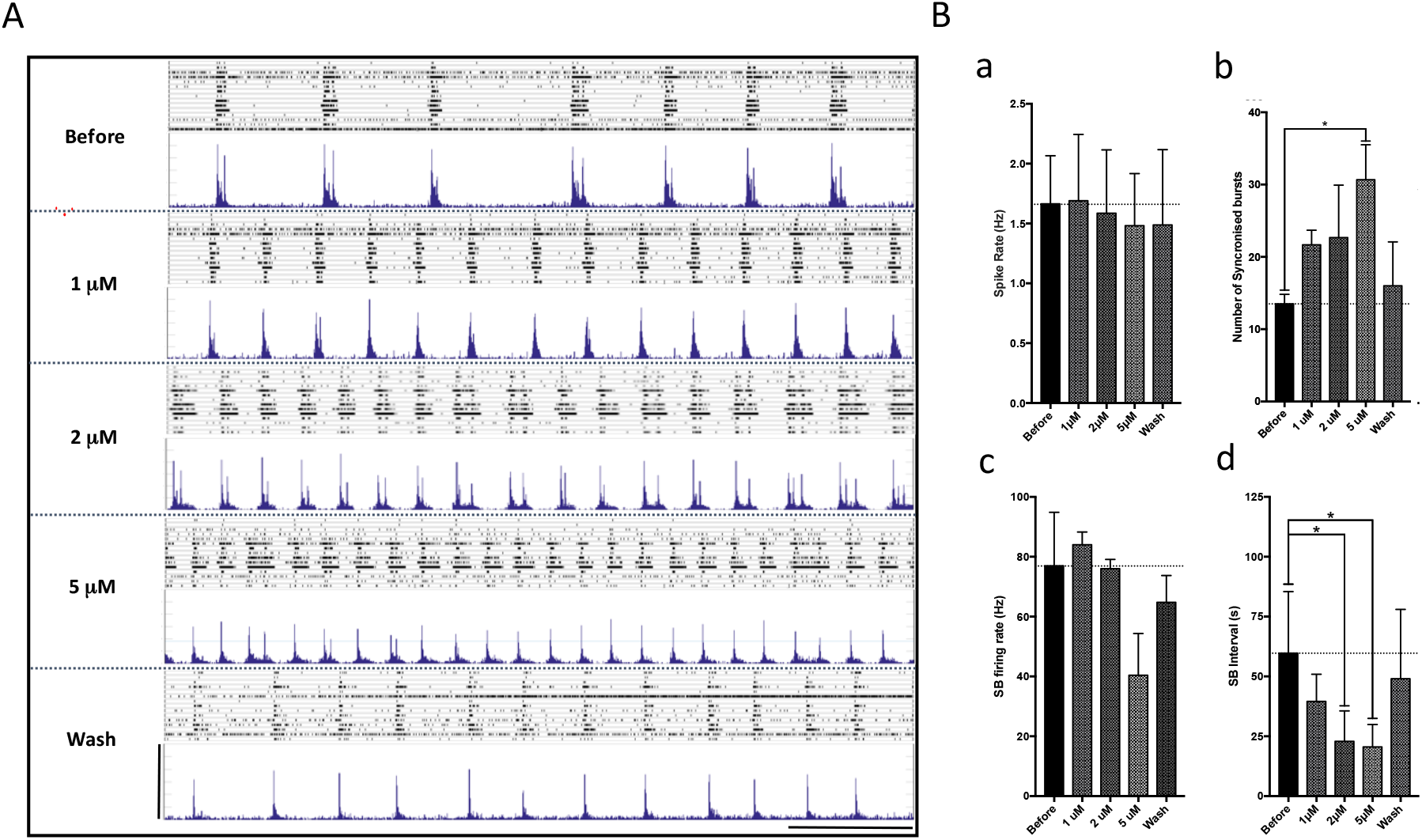
Network activity in hiPSC derived neuronal cultures is regulated by L-type voltage gated calcium channels. (A) Raster (upper panels) and ASDR plots (lower panels) of a single neuronal MEA culture before, during and after acute exposure to increasing concentrations of the L-type calcium channel blocker diltiazem. Vertical scale bar = 200 spikes per 200 ms bin; horizontal scale bar = 100 seconds. (B) Summary plots of basal and synchronised activity properties of MEA cultures in response to acute exposure to increasing doses of diltiazem: (a) Basal average spike rate; (b) average number of synchronised bursts (SBs) across the cultures; (c) Average culture-wide firing rate during SBs; (d) average interval between SBs. * = p < 0.05 as determined from Dunnett’s multiple comparisons tests following one way ANOVAs, where for Number of synchronised bursts (Bb) F_(7,21)_ = 3.324, R^2^ = 0.492 & p = 0.0107 and for SB interval (Bd) F_(7,21)_ = 6.004, R^2^ = 0.625 & p = 0.0019. All summary plots show means + SD. Number of cultures = 7.

Although the hiPSC differentiation protocol used here generates primarily cortical glutamatergic neurons, the resulting cultures also contain a small proportion on GABAergic neurons, which, in the nervous system, would be expected to exert an inhibitory effect on neural circuits and networks. To investigate the potential role of this inhibitory signalling on SBF patterns, GABA or GABA_A_ receptor antagonists were applied acutely to the MEA cultures. Exposure to 1 μM GABA caused an almost complete, but reversible, loss of both basal and hence synchronised neuronal activity (Figure 4), demonstrating that our hiPSC-derived glutamatergic neurons are GABA sensitive. In contrast, lowering the effect of endogenous GABAergic singalling within the culture by application of bicuculline, a GABA_A_ receptor blocker, significantly increased the number of SBs recorded (t_10_=2.81; p <0.05). This was accompanied by a concomitant decrease in SB firing rate (t_10_=1.91; p <0.05) and in interval length (Figure 4C; t_10_=2.14; p <0.05). As a consequence, the general spike rate was not significantly altered, so that in effect reducing the effect of GABA increased the SBF oscillation frequency without decreasing neuronal activity. These changes were reproduced when cultures were exposed to the non-competitive GABA_A_ antagonist, picrotoxin (Figure S2), strongly suggesting that this regulation is driven by the attenuation of GABAergic function.

### Synchronised network activity in MEA cultures is regulated by L-type calcium channels

Calcium signalling via L-type voltage gated calcium calcium (VGCCs) has been shown to be involved in the regulation of oscillatory behaviour in certain brain regions ^21,22^ while NMDA-mediated network signalling is thought to be regulated via calcium homeostasis mechanisms, partly through L-type VGCCs ^23,24^. Moreover, recent genetic studies have strongly implicated L-type VGCCs in increased risk for a range of neurological disorders, including schizophrenia, autism, depression and epilepsies; all pathologies with strong evidence for aberrant network signalling ^25,26^. As such, to investigate further the physiology of the neuronal cultures and to study the potential role of calcium in the underlying mechanism contributing to the network activity observed here, cells were exposed to the L-type VCCC blocker, diltiazem.

Acute exposure of increasing doses of diltiazem to the array cultures had little effect on the basal firing properties of the neurons, specifically spike rate and single-unit bursts (Figure 5A&B). However, application of diltiazem induced a dose - dependent reduction of the interval between SBs, decreasing from 62.6 ± 37.3 s, to 39.5 ± 11.4 s (1 μM), to 22.9 ± 12.7 s (2 μM) and finally to 20.5 ± 9.4 s (5 μM; Figure 5B). Dunnett’s multiple comparison tests following one way ANOVA (F=6.004, R^2^ = 0.62, p < 0.01) revealed that the reduction in SB interval reached significance at 2 and 5 μM (both p<0.05). Average SB intervals returned to baseline following washes. Up to 2 μM, diltiazem had no effect on the rate of firing within SB periods; with 5 μM exposure, the SB firing rate significantly decreased, possibly exposing the threshold at which diltiazem is acting specifically on L-type VGCCs (Figure B). The attenuation in SB intervals was replicated when cultures were exposed to the dihydropyridine-type L-type VGCC blocker nifedipine (Figure S3), strongly indicating that this observation was due to a reduction in L-type VGCC function.

Finally, to determine whether the effect on SB activity was specific to the action of L-type VGCCs or Ca_v_ channel function more generally, the Ca_v_ P/Q type blocker ω-Agotoxin TK and the Ca_v_ T-type inhibitor ML218 were also applied acutely to cultures. Exposure of the cultures to both inhibitors had no effect on the basal firing rate or, importantly, the interval between SBs (Figure 6B). Blocking both T and P/Q type channels did have an effect on the maximum rate of firing in SBs, as seen by the reduction of the SB peaks in figure 6A ASDR plots, however this did not translate to a significant reduction in SB firing rates (Figure B) at these concentrations. These results, therefore, strongly suggest that the regulation of synchronised network activity observed in these hiPSC derived neuronal cultures is due to the action of L-type VGCCs.

**Figure 6.**
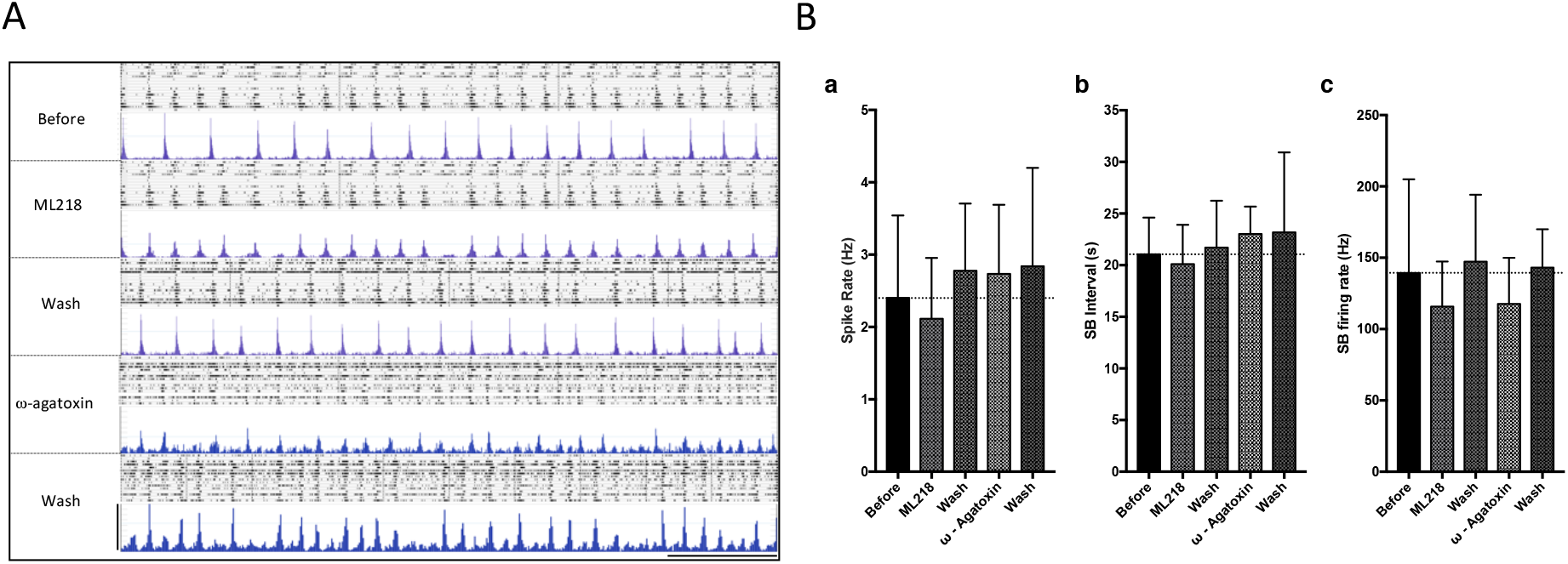
The periodicity of synchronised burst firing in hiPSC derived neuronal cultures is not affected by blocking T-type or P/Q type voltage-gated calcium channels. To determine whether the attenuation of intervals between synchronised bursts (SBs) was due to a reduction of general voltage gated calcium channel activity, neuronal cultures were exposed to the T-type calcium channel inhibitor ML218 (1 μM) and the P/Q – type blocker ω-agatoxin TK (100 nM). (A) Raster (upper panels) and ASDR (lower panels) plots showing the spontaneous activity of one MEA culture before, during and after exposure to drugs. Vertical scale bar = 200 spikes per 200 ms bin; horizontal scale bar = 100 seconds (B) Summary plots showing average (a) basal spike rate, (b) SB interval and (c) SB firing rates of profiled cultures in response to drug application. All plots show means + SD from 7 cultures.

## Discussion

We have established conditions to monitor the development of human neuronal networks derived from hiPSCs as they self-assemble *in vitro* and transition from uncoordinated, spontaneously active cultures to complex oscillatory networks with close similarities to those previously reported with dissociated rodent primary neurons. We further show that this complex network behaviour can be targeted specifically by pharmacological agents without change to the underlying basal neuronal activity. To relate this process to potential changes in network behaviour associated with mental health disorders, we demonstrated the specific effects of a pharmacological model of L-type voltage gated calcium channel hypo-function, a class of ion channels which have been strongly implicated as risk factors for psychiatric disorders.

Although the presence of synchronised array-wide activity has been reported previously in hiPSC-derived neuronal cultures ^10,27^, this reported behaviour did not progress further than the short SB firing that we observed at 40 DPP and did not evolve to the bi-stable period of oscillatory firing we report here. Indeed, this extended behaviour actually compares favourably to what has previously been described by several groups using similar MEA systems to study networks in dissociated *rodent primary neurons* ^3,5,28,29^. In particular, the extended MAP/LAP oscillations we observed in hiPSC-derived neurons at 50DPP matches to those reported in rodent studies at ~20DPP ^7,29^.

Pharmacological profiling of our cultures showed that glutamate/GABA signalling is the primary driver of the complex network behaviour observed. Whereas at initial stages spontaneous neuronal activity was insensitive to AMPA or NMDA receptor inhibition, we observed that synchronised network activity in these cultures was completely eliminated by inhibition of NMDA and AMPA receptor signalling. It has previously been suggested that blockade of NMDA but not AMPA receptors abolishes similar slow coordinated activity both *in vivo* and in *in vitro* rodent slices ^30-33^. However, work with *dissociated* rodent neurons has also shown that the coordinated burst firing is indeed blocked by both AMPA and NMDA receptor inhibition ^2,34,35^{Lu 2016). It is therefore possible that the difference seen in dissociated neurons compared to ‘intact’modelsisafunctionofthemorerandomandheterogeneousnatureofsuch cultures, which lack the intrinsic complexity, highly regulated developmental structure and region specific networking seen in slices and *in vivo*. Nonetheless, this adds to the utility of hiPSC-derived neuronal networks, as they can monitor both NMDA and AMPA receptor based glutamate signalling.

Inhibition of GABA_A_ receptor activity modifies the SBF pattern in our mature networks, decreasing the interval between SB periods, a finding consistent with reports from both rodent dissociated primary cultures ^3,7,35^ and hiPSC-derived neurons ^9,10^. This indicates that although glutamate drives the complex network behaviour, GABA shapes it. The regulation of coordinated network firing by GABAergic interneurons is well documented, ranging from coordinating oscillatory activity across brain regions ^34,40,41^, synchronised networks within structures (e.g. the hippocampus; ^42,43^) to regulation of small, localised networks ^44,45^. Furthermore, GABAergic activity is thought to be highly important throughout development, where it has been implicated in the correct formation of networks and the regulation of synaptic plasticity ^46-48^. Importantly, inappropriate GABAergic function leads to a non-physiological shift in the inhibitory/excitatory balance which is thought to underlie several neurological disorders including schizophrenia, ASD and epilepsies.

By capturing the properties of glutamate/GABA interaction of neuronal networks, the MEA neuronal networks we report here offer a cell assay platform to study the effects of risk alleles associated with neuropsychiatric disorders. To pursue this further, we examined the effects of inhibition of the L-type VGCCs Ca_v_1.2 and Ca_v_1.3, the genes encoding the core subunits of which, *CACNA1C* and *CACNA1D*, are strongly associated with neuropsychiatric disorders. The L-type VGCC blockers of diltiazem and nifedipine caused a dose-dependent reduction of the SBF interval with little change to the basal excitatory profile. This effect is specific to L-type channels and was not observed when T or P/Q channels were similarly blocked.

Ca_v_1.2 and Ca_v_1.3 alpha1subunits are expressed throughout the mammalian brain and are involved in wide array of calcium regulatory mechanisms ^49,50^. In particular, they are strongly implicated in NMDA-dependant LTP/LTD, via regulation of local intracellular calcium concentrations ^51-53^. This regulation of synaptic plasticity is thought to underlie the alterations to hippocampal – dependent learning seen with decreased or absent expression of LTCCs ^51,54,55^. Furthermore, similar rodent KO studies have implicated L-type VGCCs in hippocampal-independent mechanisms of fear learning ^56,57^ Lee 2012}^58^, regulation of axon growth ^59^, trafficking of AMPA receptor subunits ^60^ and gene expression ^61,62^.

A role for L-type VGCCs has also been described in a range of oscillatory activities in rodent neurons. Synchronised calcium transients have been shown to be mediated and controlled by L-type VGCCs in dissociated primary neurons ^[NO STYLE for: He 2006],22^, while further studies have shown that these channels can regulate calcium oscillations in intact systems ^14,15,23^. Importantly, it has also been shown that L-type VGCC currents can modulate the network response of neurons during physiological oscillatory behaviour in the hippocampus ^16,17,64^ and during modelled epileptiform activity ^65,66^. Involvement of L-type VGCCs in these systems is likely to be down to their role in the regulation of the post-burst after-hyperpolarisation (AHP), a period of hyper-polarisation which terminates high frequency firing ^67^, where they have been shown to be a key mediator of the size and duration of the AHP ^67-69^.

Our study presents for the first time the regulation of network activity in hiPSC derived neurons by modulating the function of L-type VGCCs. Importantly, this has implications for disease-modelling as mutations in the genes coding for L-type VGCCs represent some of the most strongly associated genetic risk factors for the development of mental-health disorders. Specifically, mutations in *CACNA1C* are amongst the most consistently detected genetic risk factors in both bipolar disorder and schizophrenia. Moreover, a gain-of function mutation in the pore forming region of Ca_v_1.2 is responsible for development of Timothy syndrome, a neurodevelopmental disorder characterised by cardiac arrhythmia, heart malformations and ASD ^70^.

In our study, the action of diltiazem (and nifedipine) induced a similar effect on synchronised network behaviour as that caused by inhibition of GABA_A_ receptors. Is it possible that the two pharmacological pathways share an underlying mechanism? A potential scenario is that diltiazem is acting via presynaptic L-type VGCCs on GABAergic interneurons nerve terminals. Indeed, L-type VGCCs have been shown be expressed in presynaptic terminals in the hippocampus ^71^ and in interneurons ^72^, while they have also been shown to interact with aspects of the exocytotic pathways and facilitate neurotransmitter release in certain neuronal populations ^73,74^. Furthermore, calcium currents via L-type VGCCs have been shown to act through the MAPK ERK1/2 signalling pathway, which its self has been implicated in the regulation of vesicular exocytosis ^75,76^. Importantly, this link has been observed directly as inhibition of the ERK1/2 pathway has been shown to increase neurotransmitter release via increased calcium influx via L-type VGCCs ^77^. This therefore provides a potential mechanism by which L-type VGCCs may be acting to regulate the activity of GABAergic interneurons in these iPS cell derived cultures: synchronised network activity develops in cultures after maturation of functional synapses and is dependant on both AMPA and NMDA signalling; this activity is regulated by GABAergic innervation, inhibition of which attenuates the period between high frequency culture-wide firing; blocking L-type VGCCs mimics the effect of GABA_A_ antagonism by reducing vesicular release at the interneuron-projection neuron synapse.

Our findings have shown that hiPSC derived neurons self-assemble *in vitro* to form complex, extended coordinated network firing. This has only previously been described in primary rodent cultures. This behaviour can be pharmacologically manipulated owing to the role of both intrinsic glutamatergic and GABAergic innervation and as such, serves as a physiologically relevant *in vitro* model of human neural development. Importantly, networks are sensitive to the specific attenuation of L-type VGCC function, thereby demonstrating the use of such models to monitor the function of disease relevant genetic hypo-function. The scalability of the developed MEA assays, together with increased knowledge about the genetics of mental health disorders and the increased availability of patient or engineered hiPSC lines should provide exciting opportunities for probing the function of human neural networks in disease relevant models and the potential development of novel therapeutics.

## Methods

### Cells culture and neuronal differentiation

All experiments were conducted using the IBJ4 cell line, a control iPS cell line derived from the BJ fibroblast cell line (ATCC; CRL-2522). iPS cells were differentiated into forebrain projection neurons using an adapted dual SMAD protocol ^78^. Briefly, iPS cells were maintained on matrigel (Corning) and in mTeSR1 medium (StemCell Technologies). Prior to the start of differentiation, cells were passaged onto 12 well plated coated with growth factor reduced matrigel (Corning). Once cells were >80% confluent (D0), medium was switched to N2B27 (2/3 DMEM/F12; 1/3 Neurobasal; B27-RA; N2; 1xPSG; 0.1 mM β-mecaptoethanol) + 100 nM SB and 100 nM LDN 193189. After 8 days SB and LDN were removed and cells were passaged onto fibronectin coated 12 well plates at a ratio of 2:3. After a further 8 days, cells were passaged onto PDL/laminin coated coverslips as single cells at a density of 400 cells / mm^2^. Media was switched to N2B27 with B27+ retinoic acid, and 10 μM DAPT for 7 days, after which medium was changed to astrocyte conditioned BrainPhys (Stem Cell Technologies; ^79^), supplemented with 1x B27, 1x PSG, 20 μM ascorbic acid and 20 ng/mL BDNF, in which cells remained for the remainder of the cultures. For astrocyte conditioning of BrainPhys, 50 mL of medium was added to 60-80% confluent flasks of <P5 Primary normal human astrocytes (Lonza), which were maintained previously in astrocyte medium (DMEM/F12 + 10% FBS and 1x PSG). Medium was conditioned for 72 hours at 37°C in 20% O_2_, after which conditioned medium was removed from flasks, sterilised using a 0.22 μm syringe PVDF filter and stored at at 4^°^C for use with 7 days or −20°C for longer storage. Conditioned medium was used 1:1 with fresh BrainPhys.

### Single cell electrophysiology

Patch clamping was performed on neurons using an electrophysiology rig comprising an Olympus BX51 WI upright microscope with 10x and 40x water immersion objectives; a Q-imaging Rolera Bolt CMOS camera; Luigs and Neumann Junior manipulators; Axon Instruments HS-2 Unity gain headstages; Axon Instruments Multiclamp 700B amplifier and Digidata 1550B; and a PC running Multiclamp commander. Cells were patched using borosilicate filament glass forged into pipettes using a Flaming/Brown puller (Sutter Instruments), resulting in resistances of 4-6 MΩ. All cells were patched at room temperature (around 22°C) in a basic physiological extracellular fluid (ECF) comprising 142 mM NaCl, 2.5 mM KCL, 2 mM CaCl_2_, 1 mM MgCl_2_, 10mM HEPES buffer and 30 mM D-glucose; pH adjusted to 7.4 with 4 M NaOH. The intracellular solution for pipettes consisted of 142 mM potassium gluconate, 1 mM CaCl_2_, 2 mM MgCl_2_, 10 mM HEPES and 11 mM EGTA; adjusted to pH7.4 with KOH. The osmolality of internal solutions was adjusted to 290 mOsm, using a Vapour pressure osmometer (ELITech). After braking in to cells, resting membrane potentials were determined by taking the mean voltage from 2 minutes of gap free recording (I=0). Occurrence of induced action potentials (iAPs) was determined by holding cells at around −70mV and injecting positive current steps (Δ5-10pA; 1 second) until APs were seen. If no APs were seen when the membrane potential reached −10 mV, it was assumed none would be seen at all. iAP categorisation of cells was based on visual assessment of traces. To be deemed a full AP, the overshoot had to be greater than 0 mV otherwise they were deemed as ‘attempted’. AP trains were determined as two or more full APs seen within the stimulus period. Spontaneous action potentials were observed with gap free recording with cells at resting membrane potential (I=0). Compensation for fast and slow capacitive transients was applied after seal formation and series resistance (bridge balance) values were corrected after break in and were monitored throughout experiments. Recordings were taken using Clampex 10 (Molecular Devices, USA) and were subsequently analysed using Clampfit 10 (Molecular Devices, USA).

### Multi electrode array culturing and recording

60MEA200/30iR-Ti-gr MEAs were used throughout the study (Multi channel systems; MCS); planar MEAs with 60 titanium nitrate electrodes (59 + 1 internal reference) embedded within a silicon nitrate substrate. Each electrode had a diameter of 30 μm, with 200μm spacing between electrodes. MEAs were recorded using a MEA2100-HS2×60 headstage amplifier, attached into MSC-IFB-3.0 analogue/digital interface board. Cultures were recorded using the MCRack data acquisition software running on a high-performance PC. Cultures were kept at 37°C with a TC02 temperature regulator controlled by the PC-based TCX software. The recording head stage was isolated within a custom made faraday cage.

Coverslips of cells earmarked for arrays were re-plated onto MEAs at D40-45. Clean MEAs were stored at 4°C in the dark, with the cultured area submerged in distilled water. Before cell plating, MEAs were first pre-treated with 1 ml FBS for 1 hour at 4°C in the dark, followed by washes with distilled water. Culture surfaces were then treated with 0.01% PEI (Sigma) and incubated for 1 hour at 37°C, were then washed with distilled water and left to dry completely in a sterile cell culture hood 1 hour prior to re-plating. Neurons were dissociated from coverslips with Accutase (Thermo Fisher) and plated as high density drop cultures (50,000 cells / 20 μL, which equated to around 1800 cells / mm^2^) containing 10 μg/mL laminin. After 1 hour, 1 mL of conditioned medium was then carefully flooded into each MEA and after a further 24 hours incubation, 1 ml of fresh medium was added to each array. MEA cultures were maintained in 1:1 fresh/ACM BrainPhys and in an incubator with a 2% O_2_ environment.

Raw MEA data was recorded at a sample rate of 25000 Hz and filtered online with a 200 Hz high pass and a 5000 Hz low pass filter (both 2^nd^ order butterworth). Recorded data was converted to ascii files for offline analysis using the MC Data tool (MCS). Offline analysis was achieved with custom scripts written in Matlab. Briefly, spikes were detected from filtered data using an automatic threshold-based method set at −5.5 x σ̂, where σ̂ is an estimate of the noise of each electrode based upon the median absolute deviation (MAD; ^20^). Spike timestamps were analysed to provide statistics on the general excitability of cultures. Network activity was analysed by creating array–wide spike detection rate (ASDR) plots with a bin width of 200 ms. Synchronised bursts (SBs) were detected from ASDR plots by a 4-step process: 1; A ‘start threshold’ was determined as ~ 10% of the maximum ASDR - this was varied to set the threshold just above baseline activity; 2. Data passing the start threshold was then required to pass a ‘confirmation threshold’ within 1 second – this was set as 70% of the maximum ASDR; 3. The end of SBs was determined as there being 3 seconds of < start threshold activity; 4. SBs were finalised by confirming that they lasted at least 2 seconds. Average results for every measure were calculated as medians for each culture and data from single electrodes was excluded if it contained less than 5% of the spikes detected in the most active electrode.

### Immunocytochemistry

Coverslips of cells for immunocytochemistry (ICC) were fixed as required using 4% paraformaldehyde solution (PFA; Sigma) and after washes were stored at at 4°C in the dark until use. For staining, cells were first permeabilised using 0.1% Triton X-100 (Sigma) diluted in DPBS, incubated at room temperature for 10 minutes and blocked by incubating with 1% bovine serum albumin (BSA) in PBST (DPBS + 0.1% Tween 20) for 30 mins at room temperature. Primary antibodies were diluted as required in PBST + 1% BSA incubated with cultures over-night at 4°C. Primary antibodies used in this study: mouse anti-MAP2 (Millipore, AB5622), chicken anti-TUJ1 (Neuromics, CH23005), mouse anti-GLUN1 (Antibodies Inc., 73-272), mouse anti-PSD95 (Antibodies Inc., 73-028), mouse anti-VGLUT1 (Antibodies Inc., 78-066), mouse anti-GAD67 (Millipore, MAB5406), rat anti-CTIP2 (Abcam, Ab18465) and mouse anti-SATB2 (Abcam, ab51602). The following day, cells were incubated with secondary antibodies diluted at 1/10000 in DPBS + 1% BSA for 1 hour at room temperature in the dark (Alexa Fluor 488, 594, and 647 - anti mouse, rabbit, chicken and rat as required; all Thermo Fisher) Cell nuclei were subsequently counter stained by incubating for 1 minute with 0.1 μg/ml DAPI (ThermoFisher #62248) Coverslips were mounted onto slides using DAKO fluorescence mounting medium (Agilent) and imaged using a Leica DMI6000 inverted epifluorescence microscope. Images were processed with Fiji ^80^.

### Statistical analysis

All descriptive and comparative statistics were completed with Prism 6 (Graphpad. To determine the route of analysis (parametric or non-parametric), all data was first processed using histograms, q-q plots and normality tests (kurtosis and skew tests) in R (RDevelopment, 2012). Where groups of data from the same experiment, presented with contrasting distributions, parametric tests were used as they are, in general, better equipped to cope with non-gaussian distributed data. Unless otherwise stated, all comparisons between pre- and post- treated cultures were done with paired t-tests and tests statistics, degrees of freedom and two-tailed p values are all reported. All summary plots of data show means + standard deviation. The numbers of arrays used for each experiment vary and are reported in individual figures. All results presented here were collected from at least three independent differentiations.

## Acknowledgements

Thanks to Josh Chenoweth at the Lieber Institute for Brain Development for supplying the IBJ4 iPS cell line. This work was supported by Welcome Trust Strategic award (100202/Z/12/Z), The Waterloo foundation ‘Changing Minds’ programme and AstraZeneca.

## Author Contributions

Research was designed by AJH, WP, JH, NJB and TZD. All experiments were performed by W.P. The manuscript was written and reviewed by all authors.

## Additional Information

The authors declare no competing interests.

